# Fixed Nonsynonymous vOka Vaccine Mutations in IE62 drive Varicella-Zoster Virus Attenuation in Human Skin

**DOI:** 10.64898/2026.09.28.754941

**Authors:** Reuben O. Onwe, Joao Ferreira, Megan G. Lloyd, Joseph S Flot, Paul R. Kinchington, Jennifer F. Moffat

**Affiliations:** Department of Microbiology and Immunology, SUNY Upstate Medical University, Syracuse, New York, USA; Department of Ophthalmology, University of Pittsburgh, Pittsburgh, Pennsylvania, USA; Department of Molecular Microbiology and Genetics, University of Pittsburgh, Pennsylvania, USA

## Abstract

The live, attenuated vaccine for Varicella-Zoster Virus (VZV) protects against chickenpox in children and adults. However, the vaccine strain Oka (vOka) establishes latency and may reactivate. The molecular basis of vOka attenuation remains unknown. We investigated whether three high-frequency single-nucleotide polymorphisms (SNPs) that differentiate vOka from parental strain Oka (pOka) are responsible: S628G and R958G in the immediate-early regulatory protein IE62, and *130R in the membrane protein ORF0. We evaluated how these fixed SNPs mediate the attenuated growth phenotype of VZV and diminish skin pathology by creating recombinant pOka VZV carrying these SNPs in the wild-type background. These were assessed in cells and human skin organ cultures (SOCs). In epithelial and fibroblast cell lines, the individual IE62 mutations slightly delayed viral growth at 8 – 24 hours post-infection but not in human epidermal keratinocytes (htert-HEK). The fixed ORF62 SNPs also affected ORF62 transcription, IE62 protein abundance at early times, and delayed IE62 accumulation in the cytoplasm later during infection. The *130R mutation in ORF0 conferred a growth advantage in culture, suggesting it is an adaptation to cell culture. While there was donor-to-donor variability and differences in overall virus spread in SOC, the R958G and *130R conferred an attenuated growth phenotype similar to that of vOka in SOC. Histopathologic analysis of VZV-infected skin sections revealed that, similar to vOka, S628G and R958G in IE62 limited the spread of VZV skin lesions. Our findings underscore the critical roles of conserved vOka SNPs in understanding the attenuation of the live vaccine strain vOka.

**Importance:** The live-attenuated varicella-zoster virus (VZV) Oka strain vaccine has been in clinical use for decades, yet the molecular mechanism of its attenuation remains poorly understood. Because the vaccine preparations are polyclonal and contain wild-type genotypes, they retain the ability to establish latency and reactivate, posing a risk of vaccine-induced zoster. Therefore, defining the genetic basis of its attenuation is essential for rational design of a safer, next-generation varicella vaccine. Here, we demonstrated that VZV skin lesion formation depends on the wild-type transactivator IE62, and the two fixed single-nucleotide polymorphisms, S628G and R958G, attenuate skin pathology. Specifically, we identify R958G as a key determinant of attenuated growth phenotype and skin pathology. Because R958G also serves as a diagnostic marker for vaccine strain reactivation, these findings establish a critical molecular framework for engineering safer, avirulent, and possibly highly immunogenic varicella vaccines.

## Introduction

Varicella-Zoster Virus (VZV) causes the highly contagious infectious disease varicella, also known as chickenpox [2]. During primary VZV infection, the virus establishes latency in dorsal root ganglia, from which it can reactivate to cause zoster, also known as shingles. Zoster is common in older individuals and those with immunocompromised conditions [50]. Without vaccination, zoster occurs in about 1 in 3 people and frequently causes severe long-term pain and many other neurological sequelae [35]. VZV causes 4.2 million severe complications and 4,200 deaths annually [44].

The varicella vaccine strain, vOka, was derived from a clinical isolate using standard growth-attenuation approaches involving serial passage in semi-permissive and permissive cells, namely guinea pig embryonic fibroblasts and human fibroblasts [36]. Several companies developed the vaccine, and Varivax® made by Merck and Co was approved in 1995 in the United States for universal vaccination of children and adults who had not had chickenpox. Many other countries have adopted routine varicella vaccination with related vOka strains [41]. Routine use of the well-tolerated, live, attenuated vaccine has substantially reduced varicella cases among school-age children in countries with universal vaccination programs [23, 40, 41].

The vaccine virus vOka can establish latency in recipients, and reactivation of the vaccine in some recipients can occur with virus shown to be vOka [33]. Following reports that vOka could establish latency and reactivate, sequencing the complete genome became necessary to distinguish it from the parental Oka strain (pOka). The vaccine is genetically heterogeneous, and comparative analysis across different vaccine brands (Varilrix®, Varivax®, and Suduvax®) identified 137 SNPs shared by all vOka strains [49], but with many additional manufacturer-specific SNPs being present. These analyses have revealed that most SNPs are variable mixtures of vaccine/wild type alleles, with only five SNPs showing fixation or near fixation at the vaccine allele [6, 49]. Four occur in ORF62/71, and one in ORF0 [6, 38, 49], with two in ORF62 being synonymous. Three of these fixed SNPs result in an amino acid substitution: S628G and R958G in ORF62/71 that encode the IE62 protein, and *130R in ORF0 that encodes the ORF0 protein, resulting in a lengthening of the ORF0 protein. Because five of the 54 SNPs were near fixed and specific to the vaccine, we hypothesized that these fixed SNPs drive vOka attenuation, particularly within the critical VZV protein IE62. Previous studies using the SCID-human skin xenograft mouse model showed that vOka is attenuated for replication in fetal skin compared to wild-type pOka [24] but that virus determinants of attenuation could not be identified using mixtures of vOka and pOka cosmids [51]. Transient transfection assays have suggested that vOka IE62 has less transactivation activity compared to pOka IE62, supporting roles for IE62 in vOka attenuation [5, 10, 11, 12]. The mechanisms of vOka attenuation remain unknown, and no studies have characterized the fixed, vaccine-specific SNPs in the context of viral infection.

In this study, we sought to evaluate the fixed vOka SNPs on VZV attenuation in the correct virus context using *in vitro* and *ex vivo* models. We generated vOka-like mutants with a dual reporter system that expresses luciferase and GFP from the ORF57 promoter. The viruses contained specific vOka SNPs in a cloned wild-type virus pOka BAC by recombineering. We also used a vOka clone with the dual reporter system generated by CRISPR-Cas9 gene editing. These highly related viruses allowed us to test how individual or combined fixed SNPs in IE62 and ORF0 contribute to vOka attenuation in cultured human cells and human skin, under both single-and mixed-genotype conditions.

We found that fixed IE62 mutations delayed pOka BAC growth *in vitro*, while the long form of ORF0 provided a growth advantage. The IE62 mutations reduced both its mRNA and protein levels and impaired its cytoplasmic accumulation late in infection. In human skin explants, these SNPs slightly delayed virus growth compared to pOka. However, the fixed SNPs in IE62 diminished VZV skin pathology, preventing severe blister formation. Together, these findings indicate that fixed IE62 mutations decreased the virulence and skin pathology of the vaccine virus, underscoring their critical role in attenuation.

## Material and Methods

### Viruses

The pOka bacterial artificial chromosome (BAC) used to generate mutant viruses in this study has been described previously [22, 39]. Briefly, the Tischer BAC was corrected for two spurious mutations identified by sequencing. Then it was engineered to include a reporter gene, firefly luciferase, downstream of the ORF57 coding region. The firefly luciferase gene was separated from ORF57 using a ribosomal skipping site, T2A, using BAC recombineering. The ORF57 and luciferase genes were driven by the ORF57 promoter [22]. Using the same approach, the eGFP reporter gene was inserted in frame at the same region just before the firefly luciferase gene, separated by a P2A ribosome-skipping motif. A separate manuscript describing the insertion of GFP and its use in fluorescent virus development is in preparation (JS Flot et al. in preparation; Fig.1A). To generate the mutants (Fig. 1A), we performed mutagenesis using a 2-step red recombinase scarless recombineering described previously [8]. ORF71, a duplicate of ORF62, was first deleted by recombinatorial insertion of an Ampicillin resistance cassette to replace the gene. We then developed mutations in ORF62 by site-specific recombineering. Resultant viruses restored the ORF71 gene using ORF62, and all were verified by combined short- and long-read sequencing (Plasmidosaurus Inc). All oligonucleotides used in this study were obtained from IDT, Inc. (Coralville, IA). The primer sets used are in Table S2. A Dual reporter virus derived from vOka (V1) was generated from the Varivax® vaccine using CRISPR-Cas9 gene editing. Details of how the reporter vOka was made are described elsewhere (JS Flot et al., manuscript in Preparation). All pOka BAC-derived viruses: pOka-19, S628G, R958G, S628G/R958G, and *130R were developed from transfected BACs and propagated in ARPE-19 cells. Cell stocks were made in ARPE-19 cells, which were treated with 10ug/ml Mitomycin C for four h. Viruses were frozen in aliquots in media containing 10% DMSO and 20% fetal bovine serum; all cell-associated viral stocks were titered by standard plaque assay in ARPE-19 cells and stored at -80°C.

**Figure 1:**
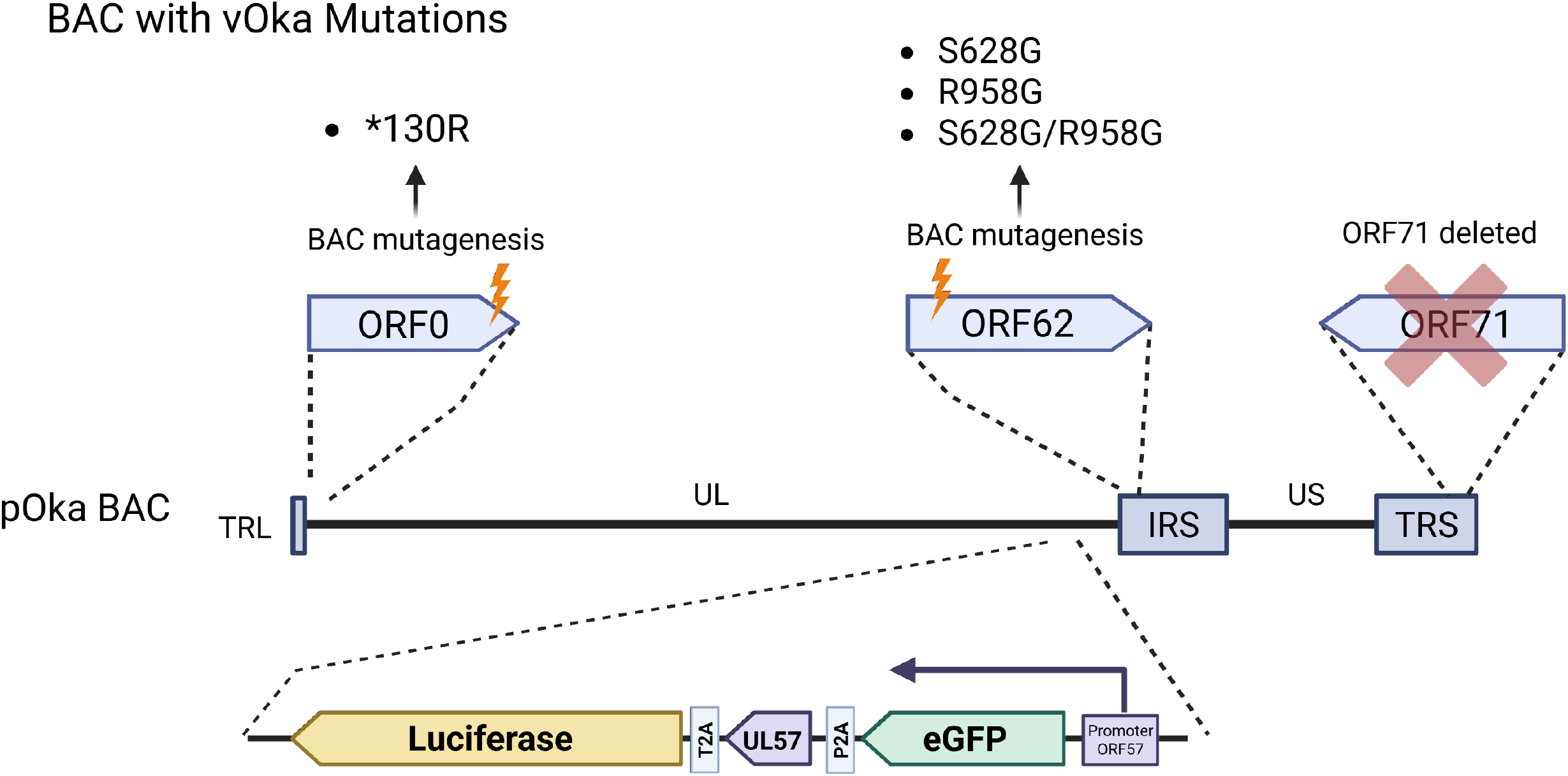
Strategy for generation of recombinant VZV containing specific vOka SNPs. (A). The center of the Figure is a schematic representation of the pOka genome that is in the pOka-19 BAC. Below this is shown the dual GFP-ORF57-l luciferase cassette in the same frame, separated by ribosome-skipping motifs T2A and P2A, as detailed in the text. The cassette was inserted downstream of the ORF57 promoter and ORF58 gene [22, 39]. ORF0 mutants were derived in this cassette. For generation of ORF62 mutations, ORF71 was first deleted by replacement with a full ampicillin resistance cassette. Details of how the mutants were made are provided in the Results and Methods section. Diagram generated using BioRender: https://app.biorender.com/illustrations/6a6b53f6d1493c01d36b1e8d?slideId=24e17b12-9dc2-47c8-8716-a3e8f1a730d7

### Cells

MRC5 cells [CCL-171; human lung fibroblast; American Type Culture Collection (ATCC), Manassas, VA, USA] were used at a passage number of less than 20. HFF cells (CCD-1137SK; human foreskin fibroblast, ATCC, Manassas, VA, USA) were used at a passage number of less than 20. ARPE-19 (CRL-2302; human retinal pigment epithelial cells, ATCC, Manassas, VA, USA) were used at a passage number of less than 40. htert-RPE (CRL-4000; human tert-immortalized retinal epithelial cells, ATCC, Manassas, VA, USA) were used at a passage number of less than 40. As described by Lloyd et al. [22], all cell types were grown in 1x DMEM (Dulbecco’s Modified Eagle Medium) with 4.5 g/l glucose, L-glutamine, and sodium pyruvate (Corning, Manassas, VA, USA). The 1x DMEM was supplemented with up to 10% heat-inactivated FBS (fetal bovine serum) (Benchmark, Gemini Bioproducts, West Sacramento, CA, USA) and an antibiotic/antifungal mix, containing penicillin/streptomycin (5000 IU/ml) and amphotericin B (250 ug/ml). MeWo cells (human melanoma cell line; kindly provided by C. Grose, University of Iowa, Iowa City) were grown as previously described [17]. Cell cultures were maintained in a humidified incubator at 37 °C with 5% CO_2_.

### Cell-free Virus Preparation

Cell-free infectious virus was prepared using a protocol slightly modified from [27]. Virus was generated using hTERT-RPE cells rather than ARPE-19 because they grow in higher density and yield substantially higher amounts of infectious VZV-infected cells and cell-free virions (data not shown). PSGC (PBS-sucrose-glutamate-serum buffer) was prepared by dissolving 5 g of D-sucrose (15621100, Thermo Scientific) in 60 mL ddH2O and 0.1 g of L-glutamic acid (BP220-1, Thermo Scientific) in 40 mL ddH2O separately, and autoclaved. The D-sucrose and L-glutamic acid were combined with 10 mL of 10× PBS and 10 mL of fetal calf serum. PSGC buffer was aliquoted and stored at −20•C. Before use, appropriate volumes of PSGC were mixed with RNase A (EN0531, Thermo Scientific) at 1:1000 and kept on ice. At least three 175-cm flasks of hTERT-RPE cells were grown to about 80% confluence and infected with cell-associated VZV at approximately 1 infected cell per 5 uninfected cells. At 80 – 90% cytopathic effect (usually day 3 post-infection), media was removed, cells were washed in PBS, and 6 mL of cold PSGC (with RNase A) was added to each flask. The infected cells were released with a rubber scraper and collected into a 50 mL Falcon tube at 4•C. Each flask was further washed with an additional 2 mL of PSGC to collect remaining cells. Cells were sonicated in an ice-water slurry bath for 3 rounds x 15s on and 15 s rest on ice to release cell-free virus, then centrifuged at 3000 × g for 45 min at 4•C. The supernatants were collected and mixed thoroughly with Lenti-X concentrator (631231, Takara) at 1:4 and kept at 4•C with gentle rocking for 30 min at 4•C. Cell-free VZV was pelleted at 1500 × g for 45 min at 4•C. The supernatants were discarded, and the pellet containing the viruses was resuspended in ∼1/50 (or ∼750 µL) of the initial volume of PSGC. The resuspended pellet containing cell-free VZV was resuspended by trituration, aliquoted (∼110 µL), and stored at −80•C or in liquid nitrogen. The cell-free virus was titered by TCID50 assay.

### Luciferase assay

MRC5, HFF, ARPE-19, htert-RPE, and MeWo cells were seeded at 6 × 10^5^ cells per well in 6-well plates. After a 24-hour incubation at 37 °C, the cell monolayers were infected with cell-associated pOka-19, vOka V1, or mutant VZV at 500 pfu per well (N = 2). At 8, 24, 48, 72, 96, and 120 hpi, infected cells were harvested, lysed, and stored at –80°C to be assayed at a later time. The luciferase assay was done according to the PROMEGA luciferase assay system (E1500; Promega) protocol and reagents. Cell lysates were combined with luciferin substrate in a black-walled glass-bottom 96-well plate and measured for luciferase flux at 570nm emission using a luminometer plate reader (Spectronics). Data were normalized to the luciferase flux at 8 hpi.

### Plaque size assay

ARPE-19, htert-RPE, HFF, MRC5, and MeWo cells were seeded at 6 × 10^5^ cells per well in 6-well plates. After a 24-hour incubation at 37•C, the cell monolayers were infected with cell-associated pOka-19, vOka V1, or mutant viruses at 100 pfu per well (N = 3). At 4 days post-infection, plates were imaged for viral plaques using an Olympus IX83 fluorescence microscope with a 10X (N.A. 0.30) objective. Images containing plaques from each virus strain, individually spaced, were acquired under the same acquisition settings with CelSens software. The fluorescent plaque area was quantified using ImageJ software, and at least 20 random plaques per virus were measured.

### RT-qPCR

ARPE-19 cells were seeded at 3 x 10^5^ cells per well in 6-well plates. After a 24-hour incubation at 37•C, the cell monolayers were infected with cell-free pOka-19, vOka V1, or mutant viruses at an MOI of 0.05; mock-infected cells served as controls. To allow viral entry, plates were incubated at 37•C for 2 h, after which the inoculum was removed. The cells were then washed with PBS, and fresh culture medium was added. At 8 and 24 hpi, RNA was harvested from infected cells using the Zymogen quick RNA/DNA miniprep kit (D7001; ZYMO RESEARCH) in accordance with the manufacturer’s protocol. RNA concentration was determined via NanoDrop and was reverse transcribed to generate cDNA using Bio-Rad iScript cDNA synthesis kit (1708891; Bio-Rad). 2 ug of the cDNA was used in a qPCR SYBR Green reaction, Bio-Rad SsoAdvanced universal SYBR Green SuperMix (1725270; Bio-Rad). For the primer sets used to amplify a specific region of the VZV ORF62, see Table S2. qPCR was run in BioRad CFX Duet system. Viral gene expression was determined by the delta Ct method.

### HEK cell culture

Human telomerase immortalized epidermal keratinocytes (htert-HEKs) were obtained from ATCC (Ker-CT, CRL 4048). These cells were grown using EpiLife CF Kit-defined HEK media (MEPICF50, Thermo Fisher Scientific), supplemented with Human keratinocyte growth supplement (S0015, Thermo Fisher Scientific) and 1% Antibiotic/Antimycotic. HEKs were grown in a low-calcium concentration (0.06 mM) to keep them in an undifferentiated state and were differentiated by switching the calcium concentration of the media to 1.2 mM using a sterile 1M CaCl^2^ solution. Cells were routinely kept at a sub-confluent density and split when density was at most 50-60% to prevent spontaneous differentiation.

### Immunoblot

Sub-confluent ARPE-19 cells were incubated at 37•C. The cell monolayers were mock-infected or with cell-free pOka-19, vOka V1, or mutant viruses at an MOI of 0.05 at 37•C for 2 h. The inoculum was removed and the cells were cultured for 8 and 24 hpi. The cells were lysed in RIPA buffer (25 mM Tris-HCl pH 7.6, 150 mM NaCl, 5 mM EDTA pH 8.0, 1% NP-40, 1% sodium deoxycholate, 0.1% SDS, and a protease inhibitor cocktail) and collected using a cell scraper. Samples were sonicated twice for 30s each. Protein concentration was determined by Bradford assay. 5 μg of protein lysate mixed with 2x Laemmli buffer (1610737, Bio-Rad) was heated at 95 °C for 5 minutes and loaded onto a pre-cast 10% SDS-PAGE gel (4568035, Bio-Rad). After electrophoresis, proteins were transferred to a PVDF membrane and blocked with blocking buffer [5% BSA in 1xTBST (Tr1is base, boric acid, EDTA, and Tween 20)]. Proteins were detected with primary antibodies prepared in blocking buffer: rabbit anti-IE62 (1:1,200 dilution, prepared in-house: see Eisfeld et al. [7] and mouse anti-GAPDH (1:3,000 dilution, GXT6274080-01, GeneTex) and secondary antibodies: goat anti-rabbit HRP (1:5,000 dilution, 31460, Invitrogen) and goat anti-mouse HRP (1:5,000 dilution, 115-035-062, Jackson ImmunoResearch). The membranes were developed with ECL substrates (170-5061, Bio-Rad) for 5 minutes at room temperature. Images were captured using a Bio-Rad ChemiDoc system, and the blots were analyzed using Fiji software (ImageJ2, Version: 2.16.0/1.54p). Densitometry of each band was determined as band intensity (integrating density divided by the area of the band) for mock, IE62 protein, and GAPDH as the loading control, and IE62 was normalized to GAPDH.

### Confocal immunofluorescence microscopy

Glass coverslips (18 mm) were placed in 12-well plates, ARPE-19 cells were seeded at confluent and incubated at 37•C. Cells were infected with cell-free pOka-19, vOka V1, and mutant viruses at an MOI of 0.3. Inoculum was removed after 2 h. At 8-and 24 hpi, cells were washed and fixed with 4% PFA in PBS for 15 minutes at room temperature. The cells were then permeabilized with 1% Triton X-100 in PBS for 30 minutes at 37 °C and blocked with blocking buffer (1% BSA and 0.5% Triton X-100 in PBS) for 1 hour at room temperature. IE62 were detected with a mouse monoclonal anti-IE62 primary antibody (1:500 dilution; Cat# GTX64189, GeneTex) and goat anti-mouse Alexa Fluor 555 secondary antibody (1:1000 dilution; Cat# A32727, Invitrogen). Coverslips were mounted onto glass slides using ProLong Diamond Antifade Mountant with DAPI (Cat# P36971, Invitrogen). Images were acquired using a Leica TCS SP8 Confocal Microscope with a 63x oil objective. Imaging settings were 1024 × 1024 resolution, 16-bit depth, and 600 Hz. Detector gain and laser intensity were optimized for each channel: blue (488) and red (555) and maintained throughout all the images acquired. Images were analyzed using both LAS X Office and Fiji (ImageJ2, Version: 2.16.0/1.54p). IE62 localization was quantified using ImageJ2. The total nuclear and cytoplasmic IE62 pixel intensity (red) was quantified by dividing the integrated density by the cell area. The nuclear integrated density was divided by the nuclear area and subtracted from the total to calculate the cytoplasmic pixel intensity.

### Human Skin Organ Culture and Viral Infection

Adult human skin organ culture was performed as previously described [9, 21]. Briefly, de-identified adult human skin, obtained from healthy, non-cancerous patients undergoing reduction mammoplasty, was thinned, dermarolled, and cut into approximately 0.3cm^2^ pieces. These skin pieces were then inoculated with 500 pfu cell-associated pOka-19, vOka V1, and mutant viruses. The skin pieces were incubated at 37 °C for 3 – 4 h and then transferred to NetWell® inserts (Corning) to establish an air-liquid interface and incubated at 35 °C. Viral spread was assessed by measuring bioluminescence (total flux, photons/sec/cm^2^/steradian) using IVIS® at 1, 3, 5, 7, 10, and 14 days post-infection. VZV cell-to-cell spread was normalized to the total flux on day 1 post-infection, and the fold change of total flux was calculated for individual skin pieces.

### Histology and immunohistochemistry

The skin organ culture model was infected with cell-associated pOka-19, vOka V1, and mutant viruses at 1000 pfu per piece, along with a mock-infection control. After inoculation, the samples were incubated at 37 °C for 3 – 4 hours to facilitate viral entry. Subsequently, infected skin pieces were transferred to NetWell® inserts to establish an air-liquid interface and incubated at 35°C. Culture media was changed every other day. At 10 days post-infection, GFP-positive areas were carefully dissected out and fixed in 4% PFA (Cat# 15710, Electron Microscopy Science). Samples were stored at 4 °C until sent to a commercial laboratory (Histowiz, Brooklyn, NY) for processing, including paraffin embedding, sectioning, and staining. Hematoxylin and eosin (H&E) staining and immunohistochemistry (IHC) with an anti-GFP antibody were used to identify VZV infection sites. Digital H&E and IHC images were returned, and images were analyzed using QuPath software version 0.6.0. Representative images are excerpts within the VZV-infected areas in the H&E and IHC digital images.

### Statistical Analysis

Statistical analysis was done using GraphPad Prism version 11 (GraphPad Prism, San Diego, CA; www.graphpad.com). Statistical significance was determined using one-way ANOVA with Dunnett’s multiple-comparison test and Student’s t-test \**p* < 0.05, \*\**p* < 0.001, \*\*\*\**p* < 0.0001.

## Results

### Generation and characterization of recombinant parental Oka with select vOka mutations

Our approach to evaluating the most fixed vOka SNPs was to introduce them into the parental Oka BAC as previously described [22, 39], and here referred to as pOka-19. We first modified the pOka BAC so that it expressed two reporter genes: eGFP, which made infected cell foci and virus in plaques easily visible and measurable. This was in addition to firefly luciferase (fLuc), driven by the ORF57 promoter and separated by the ribosomal skipping sites T2A and P2A (Fig. 1A) [22]. ORF57 is a true late, well-expressed gene that is nonessential for VZV replication [3]. ORF71, a duplicate of ORF62, was deleted so changes could be made to a single ORF62 copy in the BAC by homologous recombination in *E. coli*. When the BAC is used to derive virus, the process of virus infection reconstitutes mutated ORF71 during viral DNA synthesis and recombination in the terminal repeats, using ORF62 as a template. This was confirmed by combined long- and short-read sequencing. Recombinant viruses: pOka-19, S628G, R958G, S628G/R958G, and *130R (Fig. 1A) were recovered in ARPE-19 cells and passaged several times to restore ORF71. A different approach was used to isolate vOka with the same reporter gene cassette. To derive V1, we established an ARPE-19-based cell line expressing CRISPR-Cas9 and a guide RNA directed to the ORF57 wild-type locus. Infection of cells with Varivax® and subsequent tansfection of PCR-amplified DNA representing the dual reporter cassette resulted in GFP-positive plaques that were picked to purity (JS Flot et al., manuscript in preparation). Viruses expressing eGFP were selected and passaged 4 times to enrich for the reporter cassette and generate stocks. The virus isolated for this study was designated vOka V1. Details of its genome sequence and validations of vOka SNPs are detailed elsewhere publication in preparation (JS Flot et al.). Suffice it to state that vOka V1 has all the 5 major fixed SNPs that have been associated with vOka and approximately half of the more variable mutations now at a fixed ratio.

### Growth phenotypes in cultured cells

We assessed the growth kinetics of the mutant viruses in epithelial and fibroblast cells relevant to VZV tropism and vaccine production (Fig. 2A, B, D, E, and F). ARPE-19 (retinal epithelium), HFF (dermal fibroblast), MRC5 (lung fibroblast used for vaccine production), htert-RPE (immortalized retinal epithelium), and MeWo (melanoma) cells were infected with 500 pfu of cell-associated pOka-19, vOka V1, or mutant viruses. For the keratinocyte differentiation model (Fig. 1C), a higher virus concentration was necessary due to their higher refractility to these cells to infection. htert-HEK cells were infected with 2000 pfu of the different viruses and cultured in media containing 0.06 mM calcium. After 3 days, the media was switched to 1.2 mM calcium. This induces terminal differentiation of basal keratinocytes, which potentiates VZV replication and licenses viral late gene expression in HEK cells [13]. VZV replication kinetics were quantified by measuring luciferase activity. In general, we saw little variation in the ability to grow in all cell types. The growth of the single mutant VZV S628G and R958G was significantly reduced at 24 hpi compared to wild-type pOka-19 in the epithelial and fibroblast cells (Fig. 2A, B, D, and F; Table S1); In MRC5 cells S628G growth was reduced but not statistically significant (Fig. 2E). The pOka-19, vOka V1, and double mutant S628G/R958G strains exhibited similar growth kinetics except in htert-RPE and MRC5 cells, where vOka V1 had a growth advantage over pOka-19. This suggests that the double mutation S628G/R958G might facilitate growth in cell culture, perhaps through a compensatory mechanism. Surprisingly, the *130R mutation in ORF0, which extends the C-terminus of the ORF0 protein to 155- or 221- amino acids, enhanced VZV replication and spread in the evaluated cell lines, although it showed no difference from pOka-19 in keratinocytes. This suggests that *130R could be a cell culture adaptation. It was surprising that no differences in virus replication kinetics were observed in htert-HEK cells (Fig. 2C), which represent differentiated skin epithelium. It was possible that measuring VZV growth kinetics by luciferase enzyme activity could miss the effects of the mutations on cell-cell spread. VZV is entirely cell-associated in cultured cells, making it difficult to measure virus yield in a standard plaque assay. Thus, we evaluated cell-cell spread by infecting the epithelial and fibroblast cell lines with 100 pfu of cell-associated pOka-19, vOka V1, and mutant viruses. VZV plaque sizes were measured at 4 dpi using live-cell fluorescence microscopy (Fig. S1A–E). VZV plaques are highly variable in size and shape depending on the extent of cell fusion and morphology. In all cell lines tested, pOka-19 plaques were slightly larger than those formed by vOka V1, S628G, R958G, and S628G/R958G strains. Consistent with the growth phenotype measured by luciferase assay, the *130R strain formed slightly larger plaque sizes compared to pOka-19, vOka V1, and the ORF62 mutant strains in ARPE-19, HFF, and MRC5 cells. Similar results were obtained using cell-free virus preparations (Data not shown).

**Figure 2:**
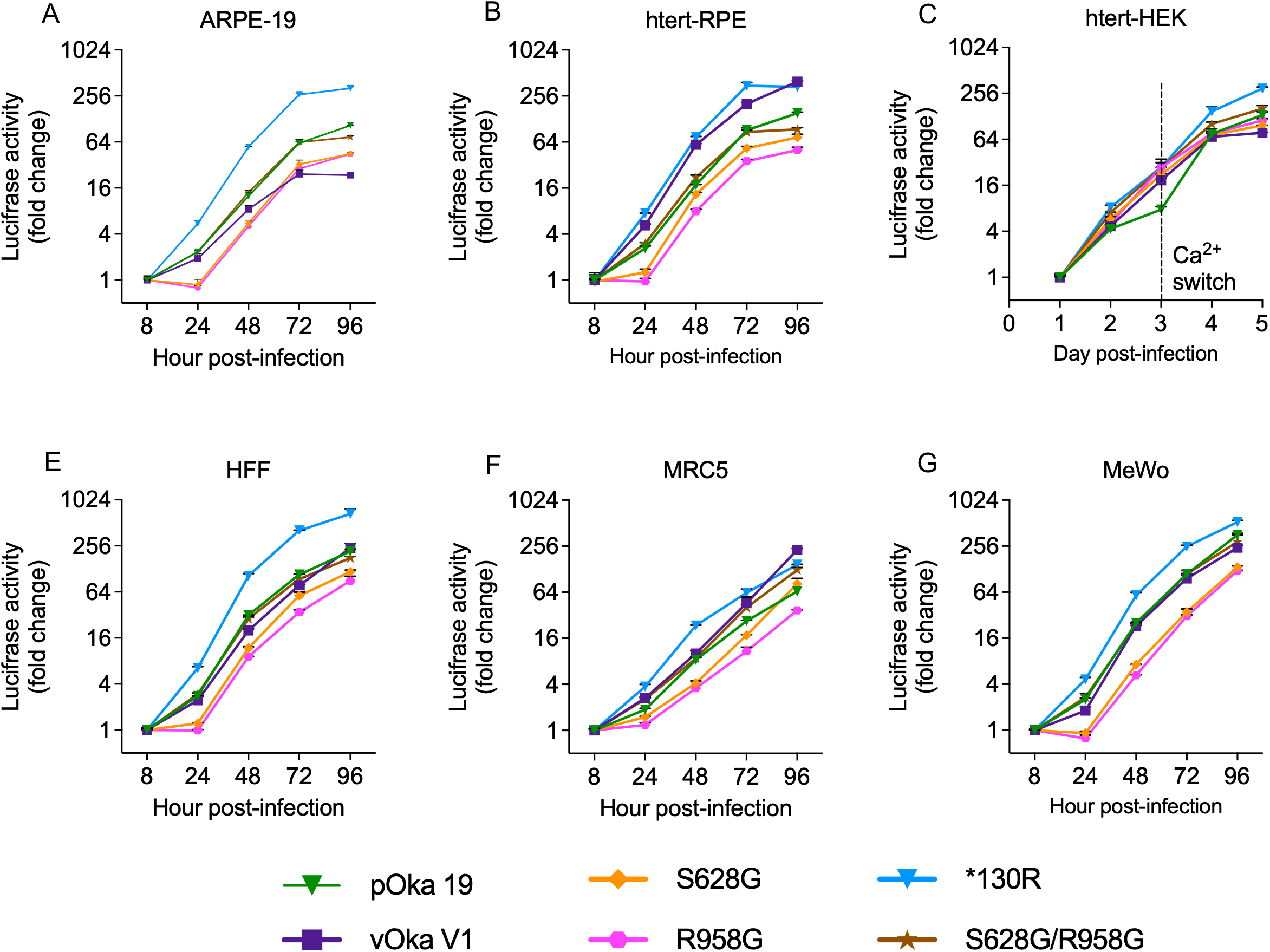
VZV growth curves initiated with low multiplicity in cultured cells (A, B, D, E and F) Epithelial (ARPE-19, htert-RPE, fibroblast (HFF, MRC5), and transformed Melanoma (MeWo) lines were infected with 500 pfu of cell-associated pOka-19, vOka V1, and mutant viruses and cultured at 37°C. (C) Htert-HEK cells were infected with 2000 pfu of cell-associated pOka-19, vOka V1, and mutant viruses, cultured at 37°C, and switched to high calcium on day 3. Viral spread was quantified by luciferase activity at the indicated time points. VZV readouts representing cell-to-cell spread were normalized to the total luciferase activity at 8 hpi, and the fold change of luciferase activity was calculated at each time point for individual viruses in each cell line. Statistical significance was determined using one-way ANOVA with Dunnett’s post hoc test for multiple comparisons. *p*< 0.05. This figure is representative of three similar independent experiments.

### Fixed vOka mutations S628G and R958G reduced IE62 mRNA and Protein in ARPE-19 cells

Because IE62 is suspected to be the major transcriptional activator protein for VZV (based on its homology to HSV-1 ICP4 and PRV ICP1), and it enhances transcription from its own promoter in plasmid transfection assays [29, 47], we investigated whether the fixed nonsynonymous mutations S628G and R958G altered ORF62/71 mRNA and IE62 protein levels. ARPE-19 cells were infected with pOka-19, vOka V1, or the mutant viruses at an MOI of 0.05 using cell-free virus. Mock-infected cells served as controls. ORF62/71 mRNA and IE62 protein levels were quantified by RT-qPCR and immunoblotting, respectively, at 8 and 24 hpi (Fig. 3). ORF62/71 mRNA levels at 8 hpi trended higher in cells infected with pOka-19 virus compared to cells infected with vOka V1, S628G, R958G, or S628G/R958G strains (Fig. 3A). By 24 hpi, ORF62/71 mRNA levels remained significantly reduced in cells infected with vOka V1 and the mutants compared to the pOka-19 (Fig. 3B). IE62 protein levels were concordant with viral mRNA. At 8 hpi, IE62 protein was significantly more abundant in pOka19 infected cells than in cells infected with the other viruses (Fig. 3C, E). In contrast, no significant differences in IE62 protein levels were detected at 24 hpi among cells infected with any of the strains (Fig. 3D, F). Taken together, these results suggest that the nonsynonymous mutations S628G and R958G may influence ORF62/71 mRNA expression and protein levels during the very early phase of infection.

**Figure 3:**
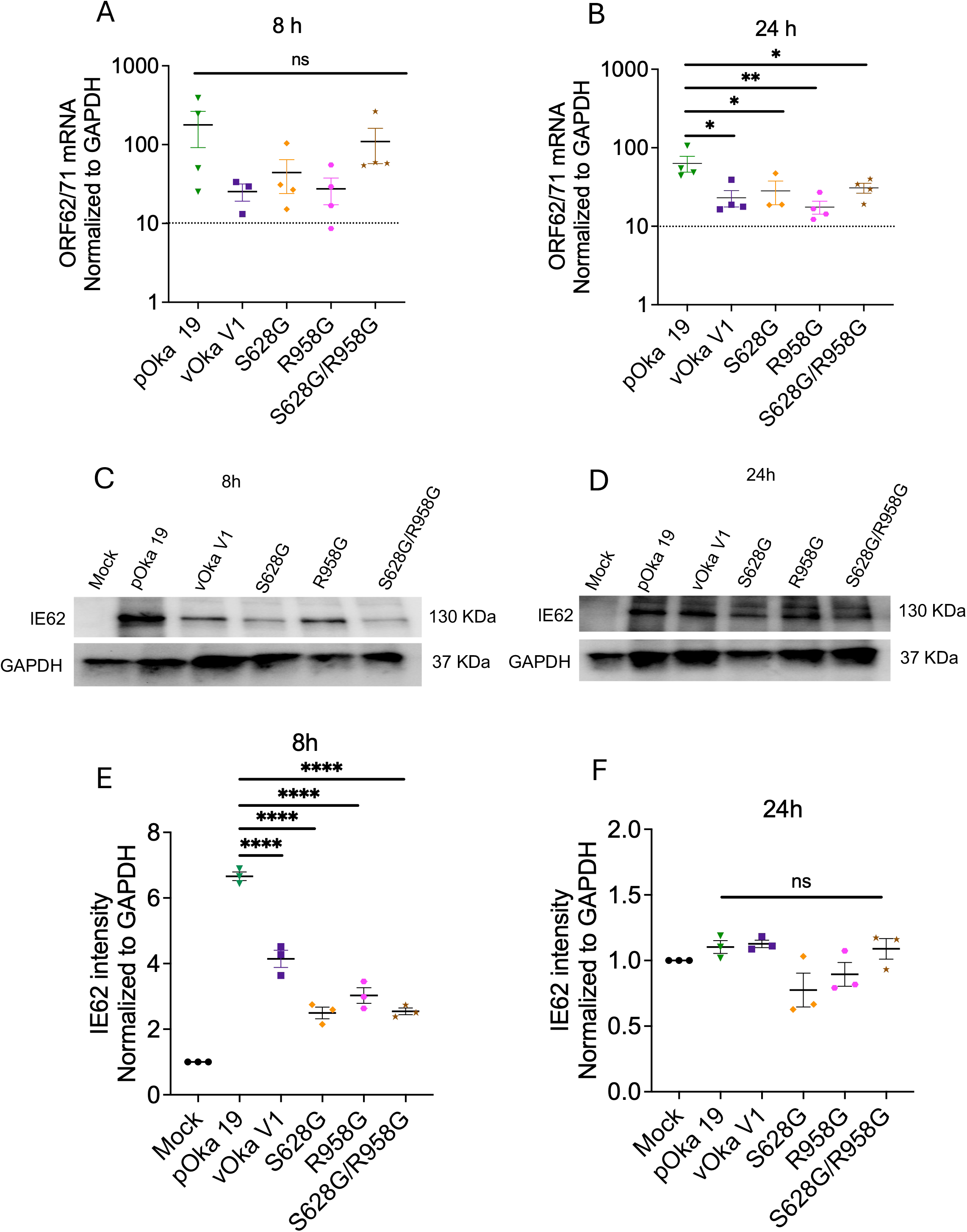
The S628G and R958G substitutions in IE62 reduced its protein levels early during infection. ARPE-19 cells were infected with pOka-19, vOkaV1, or mutant viruses at an MOI of 0.05 for 8 and 24 hpi. Mock infection served as a control. (A–B) At 8 and 24 hpi, mRNA was isolated, transcribed to cDNA, and quantified by RT-qPCR. GAPDH mRNA was quantified as a cellular control. Values are normalized to the level of GAPDH RNA. (C–D). At 8 and 24 hpi, cell lysates (5 μg) from infected cells were harvested, separated by SDS-PAGE, and transferred. Immunoblots were probed with a rabbit anti-IE62 antibody and with an antibody to GAPDH to normalize loading control. Representative blots are shown. (E–F) Densitometric quantification of IE62 levels normalized to GAPDH. Each point represents an independent biological sample (n = 3). Error bars represent SEM. Statistical significance was assessed by one-way ANOVA with Dunnett’s post hoc multiple-comparison test. \*\*\*\**p* < 0.0001.

### Fixed vOka SNPs in IE62 impact cytoplasmic accumulation

Cytoplasmic accumulation of IE62 is regulated by phosphorylation within a domain spanning amino acids 602–733, a process mediated by the viral serine/threonine kinase ORF66 [17, 18, 19]. Phosphorylation of S686 and S722 by ORF66 kinase causes IE62 to accumulate in the cytoplasm by overcoming the nuclear localization signal (residues 677-685). The fixed SNP S628G is located close to the critical regulatory domain. We investigated whether S628G and the other fixed SNPs in ORF62/71 influence IE62 protein localization by confocal immunofluorescence microscopy. ARPE-19 cells were infected with pOka-19, vOka V1, or the mutant viruses at an MOI of 0.3, and IE62 localization was assessed at 8 and 24 hpi (Fig. 4A). In pOka-19-infected cells, as expected, IE62 was found in the nucleus and cytoplasm at 8 and 24 hpi. In contrast, in vOka V1-infected cells, IE62 was almost exclusively in the nucleus at 8 hpi and was nearly undetectable in the cytoplasm. Even by 24 hpi, IE62 was in the cytoplasm of only a few vOka V1-infected cells. The mutant viruses were like vOka V1: IE62 was in the nucleus at 8 hpi and appeared in the cytoplasm in a small number of infected cells by 24 hpi. To quantify these observations, we analyzed 30 fields of view with approximately 70 cells in each field and measured the IE62 intensity (pixels per unit area) in the nucleus and cytoplasm of infected cells (Fig. 4B–E). The IE62 nuclear intensity was significantly higher in pOka-19 infected cells than in the other viruses at 8 and 24 hpi (Fig. 4B, C). The number of cells with nuclear IE62 was substantially greater in pOka-19 infected cells than in vOka V1 and the mutant viruses (Fig. 4G and H). The difference in the number of cells with cytoplasmic IE62 at 8 hpi was more pronounced, with only 5 cells infected with vOka V1 with detectable IE62 and none with the mutant viruses (Fig. 4D, E). However, the IE62 pixel intensity in the cytoplasm was similar between vOka V1 and pOka-19 at 8 hpi (Fig. 4D). By 24 hpi, IE62 cytoplasmic intensity was significantly higher in pOka-19 infected cells than vOka V1 and the mutants (Fig. 4E). Overall, fewer than 30% of vOka V1 and mutant-infected cells had any cytoplasmic IE62 signal at either 8 or 24 hpi (Fig. 4). This agrees with the RT-qPCR results showing a slight delay ORF62/71 mRNA at 8 hpi. Taken together, these results suggest that the fixed SNPs in ORF62/71 delay IE62 protein synthesis and re-localization to the cytoplasm, a process previously shown to be essential for IE62 packaging into the virion tegument during VZV assembly [1, 17].

**Figure 4:**
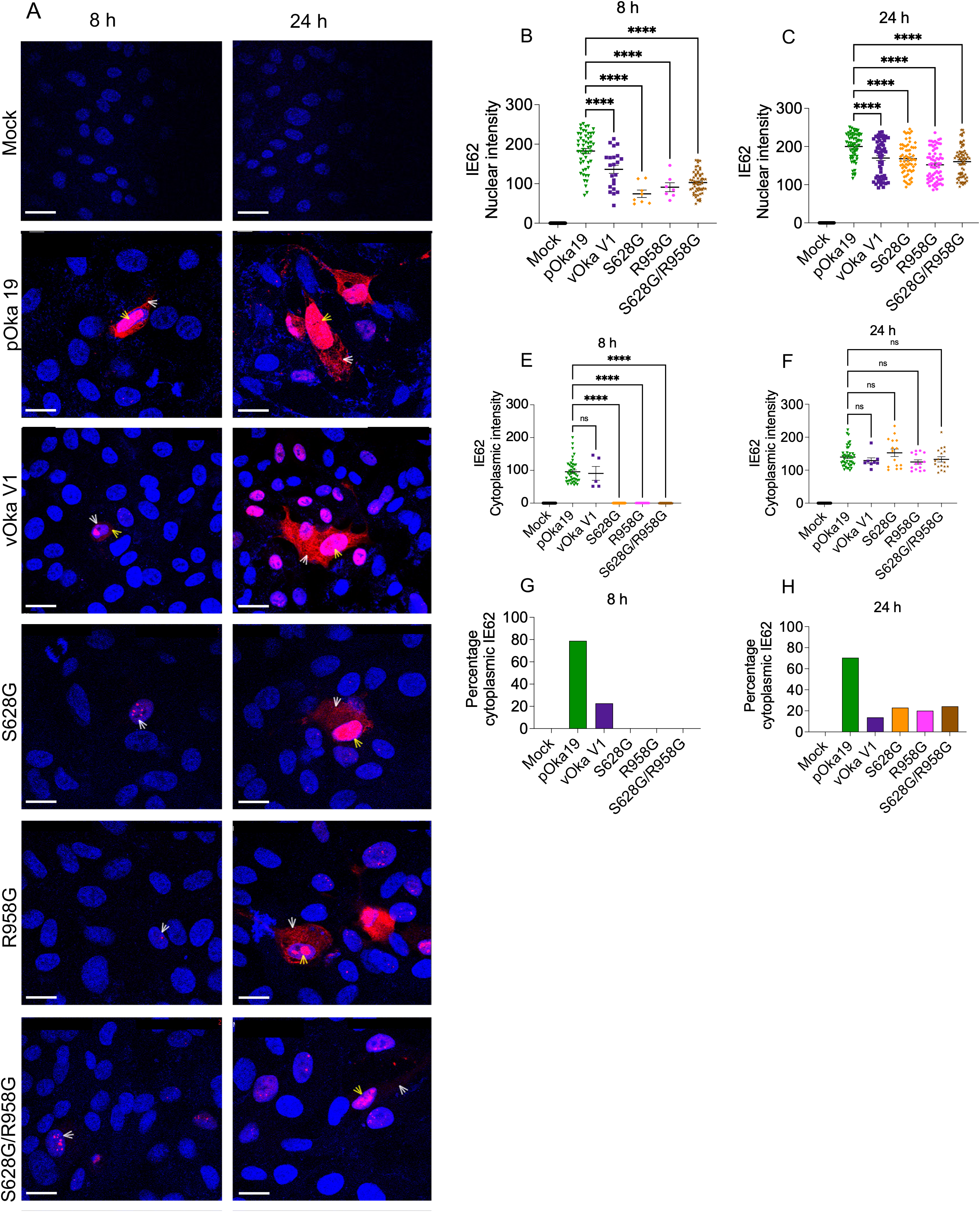
S628G and R958G vOka substitutions in IE62 delayed its cytoplasmic localization in ARPE-19 cells. ARPE-19 cells were infected with cell-free viruses at an MOI of 0.3 for 8 and 24 h. (A) Cells were fixed and stained for confocal immunofluorescence microscopy with an IE62-specific antibody (Red); nuclei were stained with DAPI (blue). The overlap is magenta. Examples of cells expressing nuclear IE62 (yellow arrowheads) and of those with cytoplasmic IE62 (white arrowheads) are indicated. Merged images are representative of 3 separate experiments. Scale bar = 50 µm. (B-E) Quantification of the confocal micrographs. For each condition, a minimum of 20 cells with positive fluorescent signals were counted and quantified by pixel intensity. Each symbol represents an infected cell, and all images were taken under the same exposure conditions. (B, C) Pixel intensity of nuclear IE62 at 8 h and 24 h. (D, E). Pixel intensity of cytoplasmic IE62 in the same cells. (F-G) Percentage of infected cells with cytoplasmic IE62. Bars are the mean ± SEM. Statistical significance was determined using one-way ANOVA with Dunnett’s post hoc multiple-comparison test. \*\*\*\**p* < 0.0001.

### Fixed vOka SNPs have delayed VZV growth phenotype in human *ex vivo* skin

The first experimental evidence of an attenuated growth phenotype for vOka was demonstrated in SCID-human skin xenografts. Compared to parental Oka and low-passage clinical isolates, vOka had significantly reduced levels of viral DNA, protein expression, and infectious virions between 14 and 28 days post-infection (DPI) [24], suggesting that the vaccine SNPs mediate vOka attenuation in human skin [34]. To investigate this further, we evaluated the related viruses using a skin organ culture (SOC) model. This model uses de-identified, non-cancerous skin obtained from reduction mammoplasty donors with informed consent [22]. Skin explants were derma-rolled and infected with 500 pfu of cell-associated pOka 19, vOka V1, or mutant viruses. Viral spread was measured by bioluminescence imaging, which is proportional to VZV-infected cells, for up to 14 days post-infection using an IVIS system. The growth phenotype in SOC was assessed separately in five skin donors to account for genetic and biological variability of the host in determining infection outcome. The combined data from three donors are presented (Fig. 5A-E). In pOka-19 infected skin, rapid viral spread was observed from 1-3 DPI, followed by a slight plateau from 5-7 DPI, and then a phase of rapid spread from 10-14 DPI that plateaued (Fig. 5A). In contrast, vOka V1 exhibited a more delayed onset of spread at 1–3 DPI, peaked at 5–7 DPI, and a decline in luciferase positivity by 12–14 DPI. Interestingly, all mutant viruses displayed a kinetic profile distinct from pOka-19 but more similar to vOka V1 during early infection, characterized by delayed spread at 1–3 DPI. However, these mutants subsequently peaked at 5 DPI, experienced a temporary plateau at 7 DPI, reached a second peak at 10 DPI, and ultimately declined by 12–14 DPI. We determined VZV infectivity in the skin pieces by calculating the percentage of pieces with viral spread at day 7 and 14 DPI (Fig. 5B). As expected, pOka-19 maintained 100% infectivity at both time points while vOka V1 and all mutant strains displayed variable but reduced infectivity at 7 DPI. More divergent phenotypes emerged by 14 DPI. The S628G and S628G/R958G mutants reached 100% infectivity, similar to the wild-type pOka-19 phenotype, whereas the R958G and *130R mutants maintained lower infectivity levels, similar to vOka V1 at 14 DPI. These suggest that vOka V1, R958G, and *130R exhibit attenuated growth phenotypes in the human skin.

**Figure 5.**
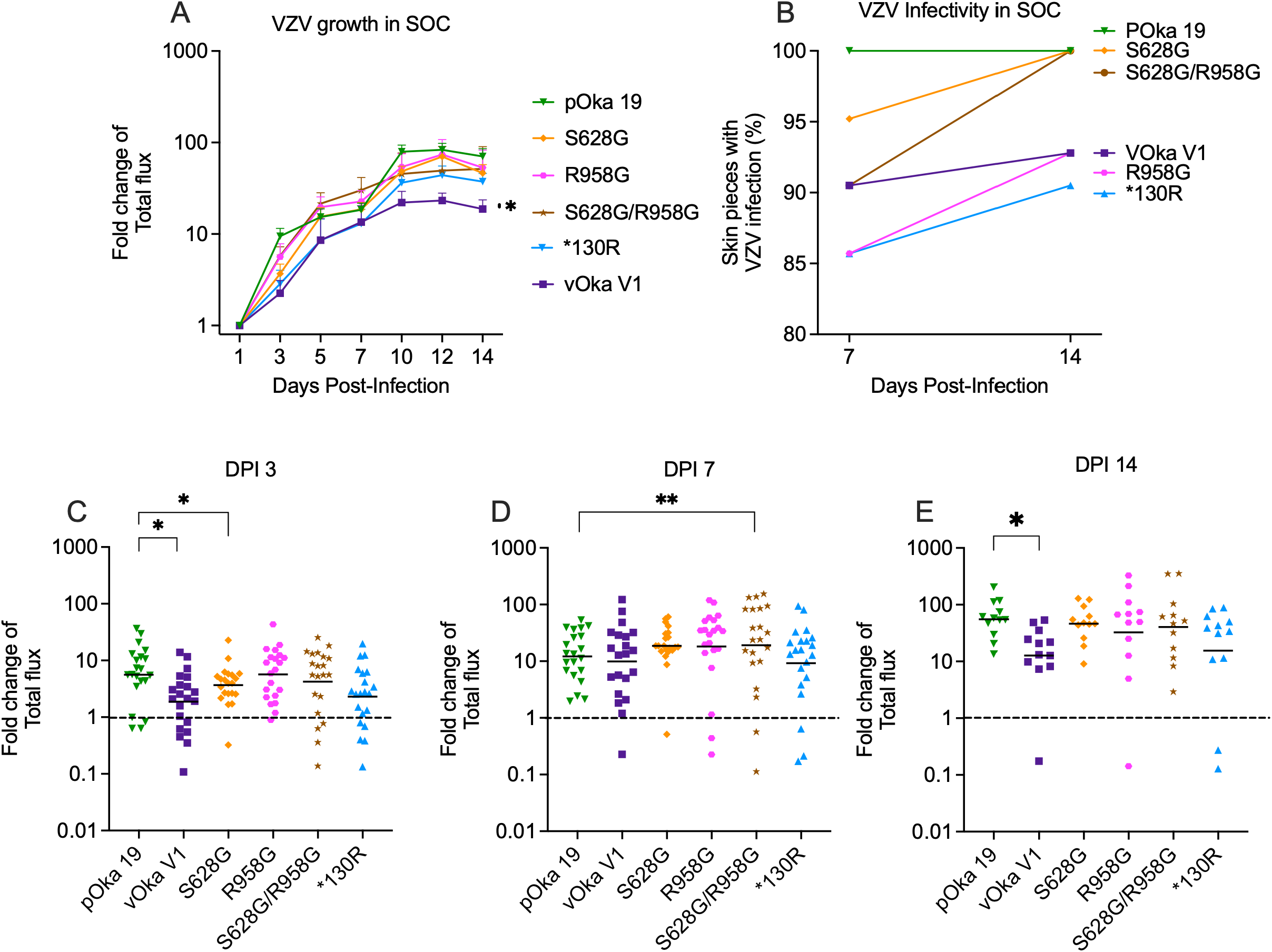
The influence of vOka SNPs on VZV Growth Phenotype in Human Skin Explants. Human adult skin pieces were infected with 500 pfu/piece with cell-associated pOka-19 vOka V1 and mutant viruses and cultured at 35°C. Viral spread was quantified by repeated daily bioluminescence imaging of the same sample (total flux, photons/sec/cm2/steradian). (A) VZV cell-to-cell spread was normalized to the total flux on DPI 1, and the fold change of total flux was calculated for individual skin pieces. Each point is the average fold change for the group (N=21); error bars are SEM. The graph includes VZV growth kinetics from three separate experiments using 3 donors. (B) Percentage of skin pieces with detectable VZV infection at 7 and 14 days. (C-E) Each point represents a single piece of VZV-infected skin (N=21), with a bar at the geometric mean at days 3, 7, and 14. A threshold was set at a fold change of 1, indicating no VZV spread. Values below the threshold (dotted lines) were excluded from the statistical analysis. Statistical significance was determined using one-way ANOVA with Dunnett’s multiple-comparison test and Student’s t-test \**p* < 0.05, \*\**p* < 0.001.

Analysis of overall viral spread per skin piece over time (Fig. S2A–F) and at specific intervals (3, 7, and 14 DPI) highlighted localized replication differences (Fig. 5C–E). At 3 DPI, viral spread was significantly lower in vOka V1 and S628G-infected tissues compared to pOka 19, an initial restriction that was not observed in the R958G mutant (Fig. 5C). By 7 DPI, the double mutant S628G/R958G exhibited significantly higher viral spread than pOka-19 (Fig. 5D). However, by 14 DPI, vOka V1 growth was significantly reduced compared to pOka 19, reinforcing its overall growth defect in skin tissue (Fig. 5E). To ensure the genomic stability of the recombinant viruses during ex vivo replication, the engineered mutations in ORF62 and ORF0 were analyzed by Sanger sequencing from viral DNA isolated post-harvest. No reversion to wild-type SNPs was detected in any of the recovered mutant strains (data not shown). Taken together, these data demonstrate that the R958G substitution in ORF62 and the *130R mutation in ORF0 serve as primary mediators of the attenuated VZV growth phenotype in human skin

### Fixed vOka SNPs attenuate VZV skin pathology *ex vivo*

Clinical observations and characterization of SCID-human skin xenografts indicated that wild-type VZV causes severe skin pathology characterized by multinucleated giant cells, ballooning degeneration, and lesions with deep dermal penetration [24, 43]. In contrast, the vOka vaccine strain causes less severe lesions with limited dermal spread [24, 51]. To determine if fixed vOka SNPs mediate attenuated VZV skin pathology, we infected skin pieces with cell-associated pOka-19, vOka V1, or mutant viruses at 1000 pfu/piece and mock infection as a control. At day 10 post-infection, GFP-positive areas were dissected for Hematoxylin and Eosin (H&E) staining. To ensure reproducibility across biological variations, we utilized samples from five independent donors, and VZV replication sites were further validated in two of the donors by immunohistochemistry using an anti-GFP antibody. Compared with mock-infected skin explants, pOka-19 and *130R mutant-infected skin showed widespread lesions with deep dermal penetration and damage to the basement membrane (Fig. 6A, B, and G). In vOka V1-infected skin, we detected viral growth more often at the edges of skin pieces, with fewer lesions and little evidence of deep dermal penetration. The basement membrane appeared intact in most sections. Surprisingly, in skin infected with VZV containing IE62 mutations, S628G, R958G, and S628G/R958G (Fig. 6C, D, E, and F), we found no lesions, few multinucleated giant cells, and an intact basement membrane. Similarly, to vOka V1 skin R958G infected the skin edges of the skin pieces, with minimal or no dermal penetration (Fig. 6E). The fixed SNPs in IE62 attenuated VZV-induced skin pathology, suggesting they are contributing factors underlying the mechanism of vOka attenuation in human skin.

**Fig. 6.**
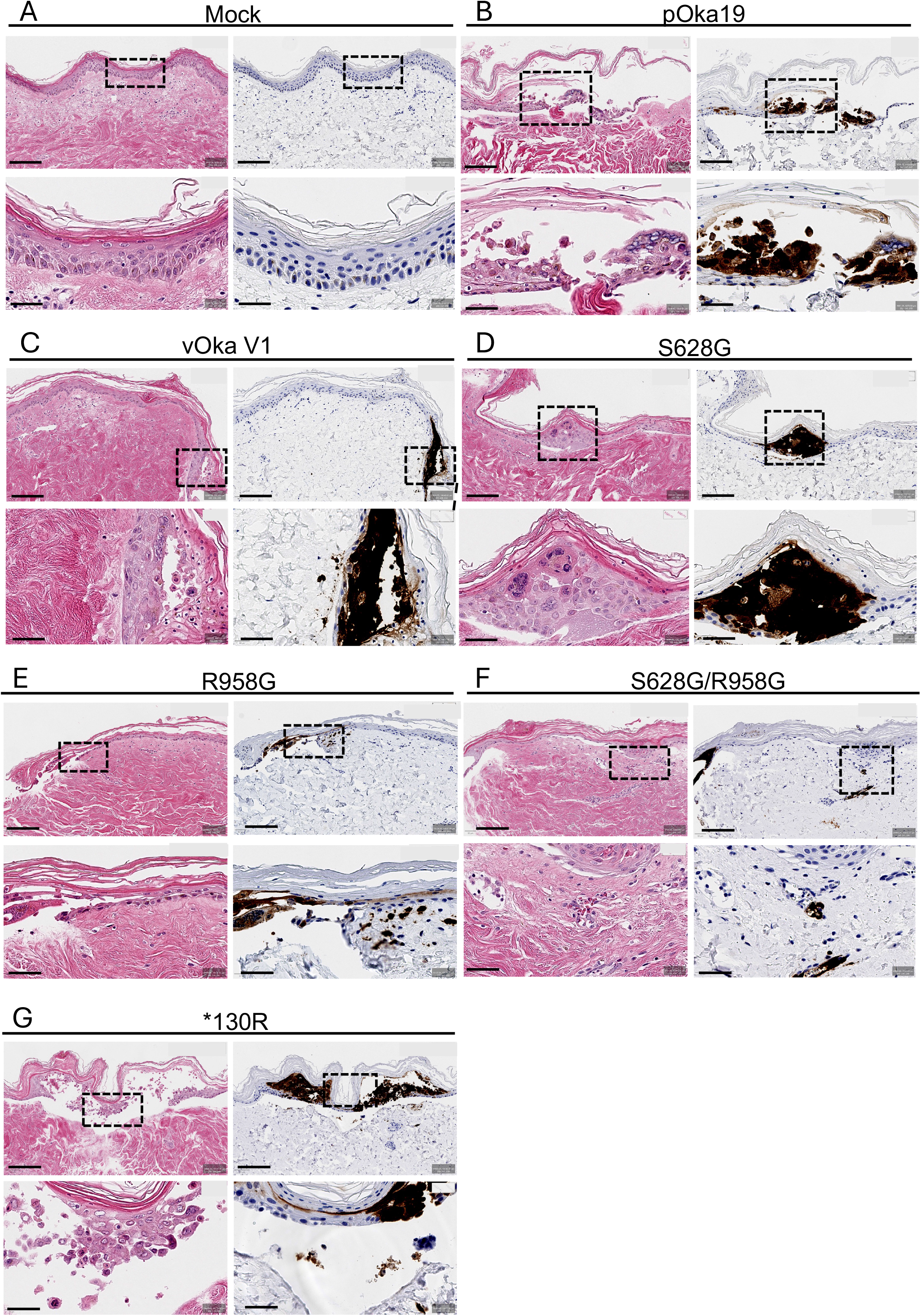
VZV-containing fixed vaccine SNPs in the IE62 protein attenuate VZV skin pathology. Skin pieces were infected with 1000 pfu/piece with cell-associated pOka 19, vOka V1, and mutant viruses, or mock-infected with uninfected cells, and then cultured at 35°C for 10 days. VZV-infected, GFP+ areas were viewed in the inverted fluorescence microscope and dissected away from uninfected areas, fixed in 4% PFA, and prepared for H&E (blue and pink) and antigen staining (dark brown) with an anti-GFP antibody. Digital slides were analyzed for histopathology and VZV-infected cells. (A-G) Representative images of mock, pOka, vOka V1, S628G, S628G/R958G, and *130R-infected skin. Scale bars: upper panels, 50 µm; and lower panels, 20 µm.

## Discussion

To date, the live attenuated varicella vaccine is the only approved human herpesvirus vaccine, which, by and large, is considered safe and effective for preventing chickenpox [48]. Zostavax®, the same attenuated strain, was also declared safe and effective at reducing zoster and subsequent postherpetic neuralgia [28]. Developed from the wild-type parental Oka using standard cell culture attenuation techniques, the live Oka vaccine strain heterogeneity has raised long-standing safety concerns regarding its clonality, molecular differences from the wild-type, and poorly defined molecular mechanism of attenuation. The early clinical trials of vOka in Japan showed that vOka reduced varicella infection [36], and vOka vaccine-associated rashes occur in less than 5% of healthy recipients [4]. The polyclonality of vOka is a major concern because most commercial vOka preparations and clinical isolates from vaccine-associated rashes retain at least 3% wild-type alleles in important ORFs [32, 34]. This has made it difficult to associate specific vOka clones with adverse events following vaccination or vaccine reactivation from latently infected neurons years later.

To address some of the challenges associated with vOka, most importantly, to elucidate the molecular mechanism of its attenuation, we created genetically defined homogeneous vOka-like VZV mutants containing fixed nonsynonymous SNPs using a pOka BAC system (Fig. 1A). We focused on the fixed nonsynonymous mutations S628G and R958G in the OR62/71 that encodes the IE62 protein, and *130R in the ORF0 that encodes the ORF0 protein. The focus on these is logical: in all of the VZV genetic amalgam, these sites are near fixed and the most frequent, and thus most likely to cause attenuation. This has been discussed previously [14, 26, 41]. IE62 is an essential multifunctional protein that transactivates its own promoter and the promoters of VZV genes of all three kinetic classes [15, 20]. Whether the fixed vaccine-specific SNPs affect any of the IE62 interactions with both the viral and host proteins is unknown. Both S628G and R958G lie in highly conserved domains thought to drive most of the protein’s critical functions, including dimerization and DNA binding for region II, while region IV likely interacts with host factors in transcriptional regulation, based on studies of HSV ICP4 [45]. ORF0 encodes the VZV glycoprotein ORF0, also known as ORFS/L. The function of ORF0 is unclear; however, its deletion results in small plaque sizes in cell culture and impaired VZV replication in the SCID-human skin xenograft model [25]. Prior investigations into vOka have relied on non-reporter vOka from the original vaccine preparation or chimeric vOka [51], which severely limits the ability to directly track vOka spread in real time. Our new dual vOka V1 overcomes this limitation using CRISPR-Cas9 gene editing from the Varivax® vaccine. The reporter vOka V1 was purified by plaque picking and contains all the major near-fixed mutations plus a binomial fraction of the vOka Variable SNPs at a near-fixed distribution of 1 or 0. The sequence of this virus is clearly vOka-like, and as we show, the virus is highly skin attenuated.

Previous studies have established that vOka has few differences or a slightly lower growth phenotype compared to pOka *in vitro* [40, 42, 51]. vOka growth was reported to be decreased in cultured keratinocytes that make up 90% of the cell types in human skin [40, 42, 46]. We found that the single mutations S628G and R958G significantly reduced VZV growth at 8 - 24 hpi compared to pOka-19 in the epithelial and fibroblast cells (Fig. 2A, B, D, and F; Table S1), but we did not detect reduced growth of vOka V1 in our keratinocyte platform. The effect of IE62 point mutations on VZV in our study is similar to the finding that mutating the serine residue at position 686 in IE62, one of the phosphorylation sites in the IE62 DBD, caused delayed VZV replication at early time points of infection in Melanoma cells [8, 15]. Interestingly, the S628G mutation is in the IE62 DBD and very close to the nuclear import signal and thus may affect those functions. However, when both mutations are present (the double mutant S628G/R958G), we found that the virus grows the same as wild-type pOka-19 and the vOka V1 across all cultured cell types tested. This could suggest that both mutations together result in a compensatory activity, and serial passage in culture may have caused them to evolve together. Passaging VZV wild-type clinical isolates extensively in culture has been shown to cause the same S628G and R958G mutations in ORF62 to arise, although not always together [16].

Additionally, highly lab-adapted strains of VZV such as VZV Ellen contain the same ORF0 mutation present in vOka [30]. We did not observe any differences in growth phenotype between the pOka 19, vOka V1, and the mutants in htert-HEK cells (Fig. 2B), suggesting that fixed SNPs do not mediate VZV attenuation in keratinocytes *in vitro*. The vOka V1 growth phenotype in htert-HEK cells is unexpected and differs from previous reports that vOka is deficient for replication in skin keratinocytes [41, 42]. However, the differences may be due to the vOka preparation used. Our study used a plaque-selected vOka V1 made from the Varivax® preparation, while the study by Tommasi et al. [42] utilized the original non-reporter Varivax® vaccine preparation. We observed that the ORF0 mutation conferred a slight growth advantage over pOka-19 in a cell type-dependent manner. This could suggest it is a culture adaptation that allowed vOka growth in non-permissive cells (guinea pig fibroblasts) during vaccine development. The plaque sizes of our VZV strains were consistent with the growth phenotype results (Fig. S1A-E).

Several previous studies have focused on IE62 transactivation activity in transfection assays and not in the context of VZV as done here. However, results were conflicting: one group reported a 68% reduction in transactivation activity for vOka IE62, whereas other groups observed only a modest decrease [31, 32]. Collectively, these findings suggest that the nonsynonymous SNPs in vOka IE62 alter its transactivation activity and, possibly, its overall biological function. Although we did not directly test whether these fixed nonsynonymous SNPs influence IE62 transactivation activity, we found that the S628G and R958G mutations reduced IE62 mRNA and protein levels in ARPE-19 cells at 8 hpi (Fig. 3A, B, C, and E). By 24 hpi, IE62 mRNA levels remained significantly reduced (Fig. 3B); in contrast, no significant differences in IE62 protein levels were detected at this later time point (Fig. 3D, F). These observations align with a previous study by Ko et al. [20], who reported that, in the absence of other VZV genes, transfecting MeWo cells with plasmids containing individual or combined nonsynonymous IE62 mutations did not alter its protein levels at 24 post-transfection. We found that the S628G and R958G mutations delayed cytoplasmic accumulation of IE62 in ARPE-19 cells at both 8 and 24 hpi (Fig. 4). Our findings align with those of Zerboni et al. [51], who demonstrated that vOka IE62 and several of their chimeric vOka IE62 constructs were predominantly nuclear at 20 hpi in MeWo cells. Consistent with their findings, the IE62 localization phenotype in cell culture is unrelated to the attenuated growth phenotype of vOka in skin [51]. Specifically, both the IE62 from our S628G mutant and the chimeric G from their study were predominantly nuclear, yet both viruses remained as infectious as pOka 19 in human skin (Fig. 5B) [51].

Using an *ex vivo* human skin organ culture developed in our laboratory to study VZV replication [21, 37], we found that these fixed vOka SNPs have a unique and divergent effect on VZV growth phenotype in human skin (Fig. 5A-B, S2A-F). The fixed SNPs caused delayed growth at early time points of infection. However, we observed that the R958G mutation in IE62 and the *130R mutation in ORF0 conferred a growth phenotype similar to the Oka V1 vaccine strain. In contrast, the S628G mutation and the double mutation S628G/R968G in IE62 resulted in a growth phenotype similar to the wild-type pOka-19 (Fig. 5B). Although the specific effect of *130R on VZV growth in human skin had not been previously investigated, Zhang et al. [52] observed an attenuated growth phenotype in SCID-human skin xenografts when the entire ORF0 was deleted. All the VZV strains in our study exhibited “hopping–like” growth kinetics believed to result from host immune control of VZV replication in the skin (Fig. 5A, S2A-F). Combined with our findings, these observations revealed that *130R and R958G mediate an attenuated growth phenotype in human skin.

Lastly, the clinical hallmark of both varicella and zoster is a distinctive vesicular rash. The pOka skin pathology is usually characterized by ballooning degeneration and lesions that penetrate deep into the dermis from the epidermis, during which the basement membrane is degraded [37]. We found, using H&E staining paired with immunohistochemistry of GFP-positive sections, that tissues infected with pOka-19 and the *130R mutant exhibited severe lesions (Fig. 6A and E). These pathologic features were highly consistent with the classic wild-type VZV skin pathology described by Moffat et al. [24] and Taylor & Moffat [37]. In contrast, the vOka V1, S628G, R958G, and S628G/R958G variants did not cause severe skin pathology (Fig. 6B, C, and D) as there were fewer severe vesicles, and/or no vesicles, little ballooning degeneration, and an intact basement membrane in skin sections infected with these viruses. These results provide the first direct evidence defining the role of fixed vOka SNPs in modulating VZV skin pathology. Specifically, our findings demonstrate that fixed nonsynonymous mutations in the IE62 act as a key determinant that attenuates vOka skin pathology.

Generally, we found that the fixed vOka SNPs in IE62 delayed VZV growth both in vitro and ex vivo, reduced IE62 mRNA and protein levels, and delayed the cytoplasmic accumulation of IE62 at early times of infection (Fig. 7). Both the vOka V1 and the mutant viruses exhibited delayed growth in *ex vivo* infection. Most importantly, the fixed vOka SNPs in IE62 attenuated VZV skin pathology, demonstrating that the fixed nonsynonymous vaccine SNPs in the vOka IE62 protein form the genetic basis for vOka attenuation.

**Figure 7:**
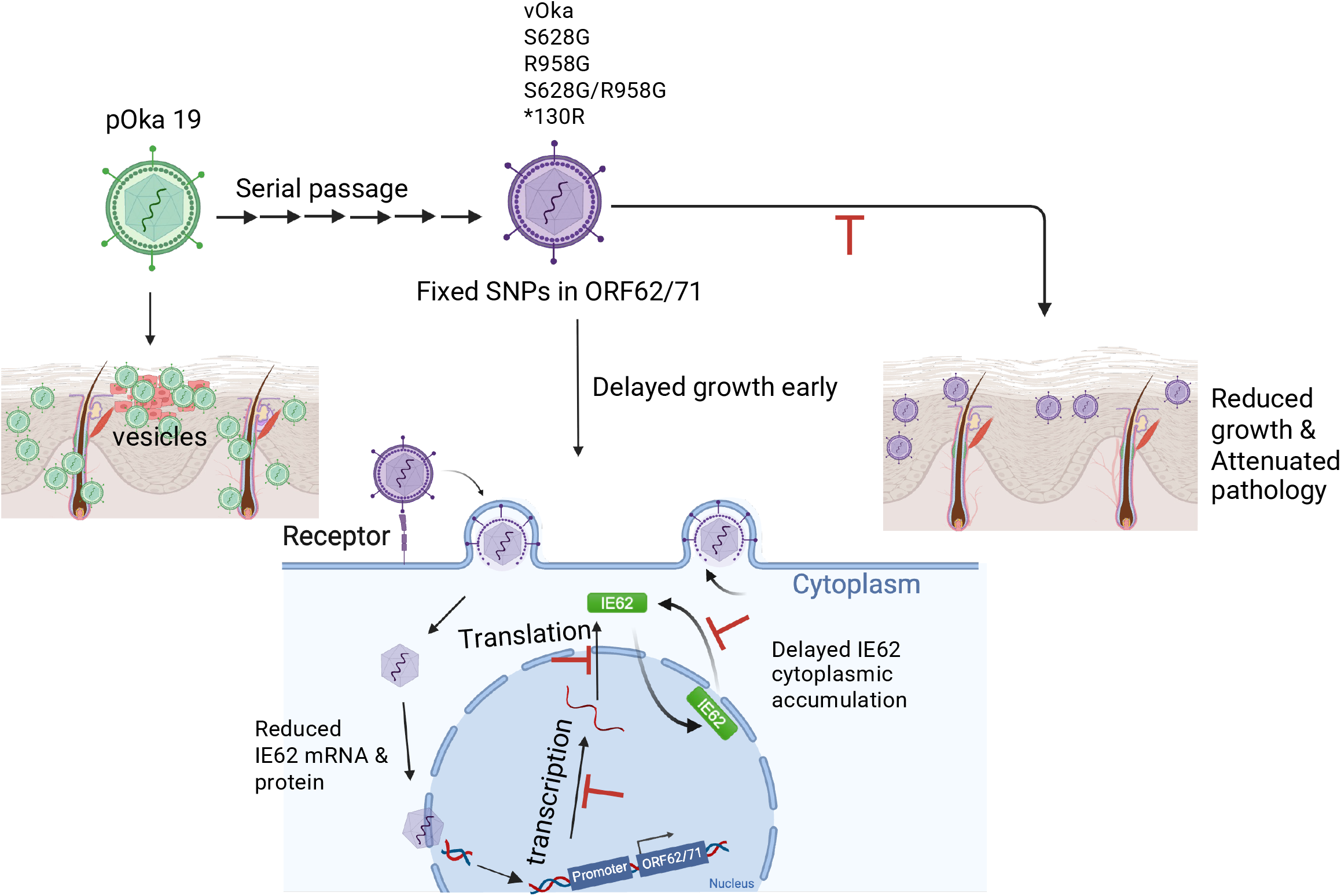
The fixed mutations in IE62 mediate attenuation of the Oka vaccine strain in human skin. Our data suggest that the fixed nonsynonymous mutations in IE62 delayed growth in epithelial and fibroblast cells as well as skin organ culture at early time points of infection. They also reduce levels of IE62 mRNA, protein, and cytoplasmic accumulation at an early time point of infection. These fixed IE62 SNPs attenuated VZV skin pathology. The model was created in BioRender: <u>tps://app.biorender.com/illustrations/6a6b55f50fb06fc53792f271?slideId=5911e21f-561d-4040-8679-cb7143d40483</u>

## Figures Legends

Fig. S1 VZV plaques reflect the effect of the fixed vOka SNPs on growth phenotype in cell culture. ARPE-19, htert-RPE, HFF, MRC5, and MeWo were infected with cell-associated pOka-19, vOka V1, and the mutant strains. At 4 days post-infection, images of plaques were acquired by a live fluorescence microscope. (A-E) Areas of at least 20 plaques were measured for infected cells in each of the VZV strains. Each symbol represents a plaque. Statistical significance was determined using one-way ANOVA with Dunnett’s multiple-comparison test and Student’s t-test. \**p* < 0.05, \*\**p* < 0.001.

Fig. S2. VZV Growth Phenotype in individual Human Skin pieces. Human adult skin explants were infected with 500 pfu per piece of cell-associated pOka-19, vOka V1, and mutant viruses and cultured at 35°C. Viral spread was quantified daily by bioluminescence (total flux, photons/sec/cm^2^) using an IVIS imaging system at the indicated time points. The growth phenotype of individual viruses in SOC from the combined three independent donors (n = 21 biological replicates) is shown: (A) pOka, (B) vOka, (C) S628G, (D) R958G, (E) S628G/R958G, (F) *130R.

## Data availability

All the data and reagents described for this paper will be made available upon request.

## Acknowledgements

This work was supported by a Public Health Service award from NIH (NIAID R01 AI158510). The authors acknowledge contributions by studies performed under P30 EY08098 and Unrestricted awards to the Department of Ophthalmology, University of Pittsburgh by the Pittsburgh Eye & Ear Foundation and Research to Prevent Blindness Inc., NY. JRF was a T32 predoctoral Recipient under T32 AI049820-20 (N DeLuca PI)

